# LINE-1 ORF1p RIP-seq reveals widespread association with p-body enriched mRNAs

**DOI:** 10.1101/2020.08.20.259424

**Authors:** Erica M. Briggs, Wilson McKerrow, Paolo Mita, Jef D. Boeke, Susan K. Logan, David Fenyö

## Abstract

**Background:** Long INterspersed Element-1 (LINE-1) is an autonomous retroelement able to “copy-and-paste” itself into new loci of the host genome through a process called retrotransposition. The LINE-1 bicistronic mRNA codes for two proteins, ORF1p, a nucleic acid chaperone, and ORF2p, a protein with endonuclease and reverse transcriptase activity. Both proteins bind LINE-1 mRNA in cis and are necessary for retrotransposition. While LINE-1 transcription is usually repressed in most healthy somatic cells through a plethora of mechanisms, ORF1p expression has been observed in nearly 50% of tumors, and new LINE-1 insertions have been documented in a similar fraction of tumors, including prostate cancer.

**Results:** Here, we utilized RNA ImmunoPrecipitation (RIP) and the L1EM analysis software to identify ORF1p bound RNA in prostate cancer cells. We identified LINE-1 loci that were expressed in androgen sensitive and androgen independent cells, that we show are representative of LINE-1 copies expressed in prostate cancer before and after treatment. In all androgen independent cells, we found higher levels of LINE-1 RNA, as well as unique expression patterns of LINE-1 loci. Interestingly, we observed that ORF1p bound many non-LINE-1 mRNA in all prostate cancer cell lines evaluated, and polyA RNA, and RNA localized in p-bodies were especially enriched. Furthermore, the expression levels of many of the identified ORF1p bound mRNAs also correlated with expression of LINE-1 RNA in prostate tumors from The Cancer Genome Atlas (TCGA).

**Conclusion:** Our results show a significant remodeling of LINE-1 loci expression in androgen independent cell lines when compared to parental androgen dependent cells, suggesting an evolution of LINE-1 expression during prostate cancer progression. Additionally, our finding that ORF1p bound a significant amount of non-LINE-1 mRNA, and that the enriched ORF1p bound mRNAs are also amplified in LINE-1 expressing TCGA prostate tumors, suggest that ORF1p may play a role in non-LINE-1 RNA processing and regulation of specific transcripts in prostate tumors.

## Introduction

Many factors have played a role in shaping the evolution of the human genome. In particular, the Long INterspersed Element 1 (LINE-1) retrotransposon, has significantly contributed in shaping the size, structure, and expression of the human genome over millions of years [1, 2]. Since LINE-1 mobilizes through retrotransposition, a copy and paste mechanism that utilizes an RNA intermediate, LINE-1 sequences have accumulated in the genome of virtually all organisms, and today over 17% of the modern human DNA is comprised of LINE-1 copies [3]. An estimated 500,000 copies of LINE-1 exist in the human genome, however, the majority of these sequences are unable to mobilize due to truncations or incapacitating mutations and inversions. However, an estimated 80-100 full length LINE-1 sequences, belonging to the subfamily L1Hs, are retrotransposition competent. Among these full-length LINE-1s, only a few “hot” loci contribute to the bulk of LINE-1 mRNA expression [4, 5].

Full length LINE-1 sequences consist of a 5’UTR/promoter, two open reading frames, coding for ORF1p and ORF2p proteins, and a 3’ UTR with a polyA signal [6]. Following transcription by RNA polymerase II, LINE-1 mRNA is exported from the nucleus. ORF1p and ORF2p are translated in the cytoplasm, where they bind LINE-1 mRNA and form LINE-1 ribonucleoproteins (RNP) composed of LINE-1 mRNA coated by many ORF1p trimers and presumably one or a few ORF2p. In dividing cells, the LINE-1 RNPs can enter the nucleus upon breakdown of the nuclear membrane during mitosis [7]. ORF2p then nicks the DNA in A/T rich regions (AA/TTTT consensus) using its endonuclease domain, and inserts a new copy of LINE-1 through its reverse transcriptase domain [8, 9]. While LINE-1 has demonstrated strong cis preference in mobilizing its own mRNA, its proteins have also been shown to bind and mobilize non-LINE-1 mRNA such as SINEs and other mRNAs that produced processed pseudogenes [10–12].

Studies have demonstrated that both LINE-1 proteins, ORF1p and ORF2p, may be necessary for LINE-1 retrotransposition [13]. ORF1p is a nucleic acid chaperone and is composed of a coiled coil domain, an RNA recognition motif (RRM), a carboxy-terminal domain (CTD) and an unstructured N-terminal region [14, 15]. The coiled coil domain is responsible for the formation of ORF1p homotrimers. The RRM, CTD, and ORF1p trimerization facilitates ORF1p binding to single stranded nucleic acids [14]. In studies of LINE-1 overexpression, ORF1p has been shown to have a strong cis preference to LINE-1 mRNA [16]. In addition to binding LINE-1 mRNA, ORF1p has also been shown to bind ssDNA. ORF1p has also been shown to aid in strand exchange and annealing, suggesting a possible role in directly facilitating LINE-1 reverse transcription [17, 18]. While ORF1p seems to be needed for LINE-1 and pseudogene retrotransposition, it is not crucial for SINE retrotransposition [10] although, increased levels of ORF1p were shown to promote higher SINE retrotransposition [19]. ORF1p predominantly localizes to the cytoplasm, and has been found in stress granules (SG) and in close proximity to processing bodies (p-bodies) [20, 21]. Stress granules and processing bodies are both cytoplasmic RNA granules that play a role in RNA metabolism and translation regulation. Stress granules form in response to cellular stressors such as heat shock and oxidative stress, and have been shown to accumulate mRNA and proteins that are stalled during translation initiation. Processing bodies on the other hand are present in non-stressed cells and contain machinery involved in RNA decay, RNA mediated gene silencing, RNA storage, and translational repression. [22–24] Since ORF1p can be translated with high efficiency and specific antibodies are commercially available, its endogenous expression and localization have been widely studied. In contrast, ORF2p expression is strictly regulated due to an unconventional translation, leading to almost undetectable endogenous levels [25, 26]. Yet, evidence of ORF2p endonuclease and reverse transcriptase activity is clear through its impact on genome evolution and, as more current findings show, through the identification of many new LINE-1 insertions in several cancers [13, 27, 28].

DNA methylation, histone modifications, and RNA interference all limit LINE-1 expression and function in somatic cells [29–32]. However, mechanisms that limit LINE-1 expression are often dysfunctional in cancers, allowing for the expression and mobilization of LINE-1 [28, 33, 34]. LINE-1 expression has the potential to disrupt genomic stability, making it a likely component in cancer progression [35, 36]. LINE-1 ORF1p expression has been observed in around 47% of tumors tested, and roughly in 40% of prostate tumors [33]. In some cancers, such as pancreatic ductal adenocarcinomas and breast cancer, ORF1p expression patterns correlated with poorer clinical outcome [37, 38]. New LINE-1 insertions have been identified in 53% of tumors sequenced, including around 60% of prostate tumors. Additionally, the LINE-1 insertion rate in prostate tumors was higher in metastatic tumors when compared to primary tumors, suggesting a correlation between LINE-1 retrotransposition and tumor progression [28].

The repetitiveness of LINE-1 sequences poses a challenge to identifying actively transcribed LINE-1 loci [39, 40]. However, a newly developed analysis software, LINE-1 Expectation Maximization (L1EM), utilizes the expectation maximization algorithm to overcome this obstacle and identify loci of actively transcribed LINE-1 [41]. Here, we performed RNA-immunoprecipitation of ORF1p to identify ORF1p bound transcripts and utilized L1EM to identify specific LINE-1 loci expressed in androgen sensitive prostate cancer cells, and androgen independent clones. Among the cell lines tested, we found a high degree of variation in the specific loci expressed, with LNCaP derived cell lines, LNCaP-95 and LNCaP-abl, having LINE-1 expression patterns that are more similar to each other and to LNCaP than to an unrelated prostate cancer cell line (22Rv1). Surprisingly, we found ORF1p bound significant levels of non-LINE-1 mRNA, including enrichments of circRNA, polyadenylated RNAs, and RNAs associated with p-bodies. Interestingly, we show that the specific expression of the non-LINE-1 RNAs enriched in the ORF1p immunoprecipitations correlates with LINE-1 mRNA expression in TCGA prostate cancer samples, suggesting a possible role of ORF1p in RNA accumulation and processing of these specific transcripts.

## Results

### LINE-1 loci expressed in prostate cancer cell lines

We recently showed that our tool, L1EM, is able to quantify RNA expression at specific LINE-1 loci from RNA-seq data [41]. We also found that accurately predicting this expression requires significantly deeper coverage compared to a standard RNA-seq analysis. Here, we performed ORF1p RNA immunoprecipitation (RIP-seq) in four prostate cancer cell lines (LNCaP, LNCaP-95, LNCaP-abl and 22Rv1) to enrich for LINE-1 reads and more accurately measure the expressed loci (Figure 1A). LNCaP cells are representative of an earlier stage, androgen sensitive prostate cancer, while its androgen independent clones, LNCaP-95 and LNCaP-abl, are more representative of treatment resistant prostate cancer, a later stage in disease progression. These cell lines were used to address the question of whether the LINE-1 expressed loci would change with selective pressure on the parental LNCaP cell line to evolve into an androgen independent sub-clone (LNCaP-95 and LNCaP-abl). 22Rv1 cells are also an androgen independent prostate cancer cell line, but are not clonally related to the LNCaP cell line. While LINE-1 comprises a substantial percentage of the human genome, many loci contain mutations resulting in truncated LINE-1 mRNA. We found strong enrichment for LINE-1 RNAs in all cell lines, confirming our previous results [21]. As expected, the greatest enrichment was measured for loci that retain a full length ORF1 compared to those that have acquired an ORF1 nonsense mutation (Figure 2A). Enrichment was even greater at loci that also have a full length ORF2. Given that ORF1p is assumed to have a cis-preference, this result validates our ability to immunoprecipitate ORF1p bound to RNA and to assign the immunoprecipitated LINE-1 mRNA to specific genomic loci. Lower enrichment of LINE-1 mRNA in the androgen independent cell lines (LNCaP-95, LNCaP-abl and 22Rv1) is primarily due to higher levels of LINE-1 in the input RNA (LINE-1 quantifications before and after ORF1p-IP are shown in Figure S1). We then looked at the specific intact (full-length and containing no nonsense mutation) LINE-1 loci that are expressed in each cell line. Figure 2B shows a heat map of the ten most highly expressed intact LINE-1 loci; an extended heatmap with 25 loci is provided in Figure S2, and the full list of LINE-1 loci can be found in table S1. Overall, we find that the expressed loci differ widely between the cell lines. The 22q12.1 locus is highly expressed in most of the cell lines, but is not detected in LNCaP-95 in which the locus at 8q24.21 is most highly expressed, but not detected in any other cell line. Overall, the LINE-1 expression pattern is more different than similar between cell lines, but has greater overlap among the LNCaP and LNCaP-derived cell lines than between the LNCaP “family” cell lines and 22Rv1. The overlap in expressed LINE-1 loci ranges from 35% overlap between LNCaP and LNCaP-95 to 42% between LNCaP and LNCaP-abl compared to a 6% overlap between LNCaP-95 and 22Rv1 and 28% for LNCaP and 22Rv1 (one-sided t-test p=0.03). This likely reflects the fact that LNCaP-95 and LNCaP-abl are derived from LNCaP through androgen deprivation, whereas 22Rv1 is clonally independent.

**Figure 1.**
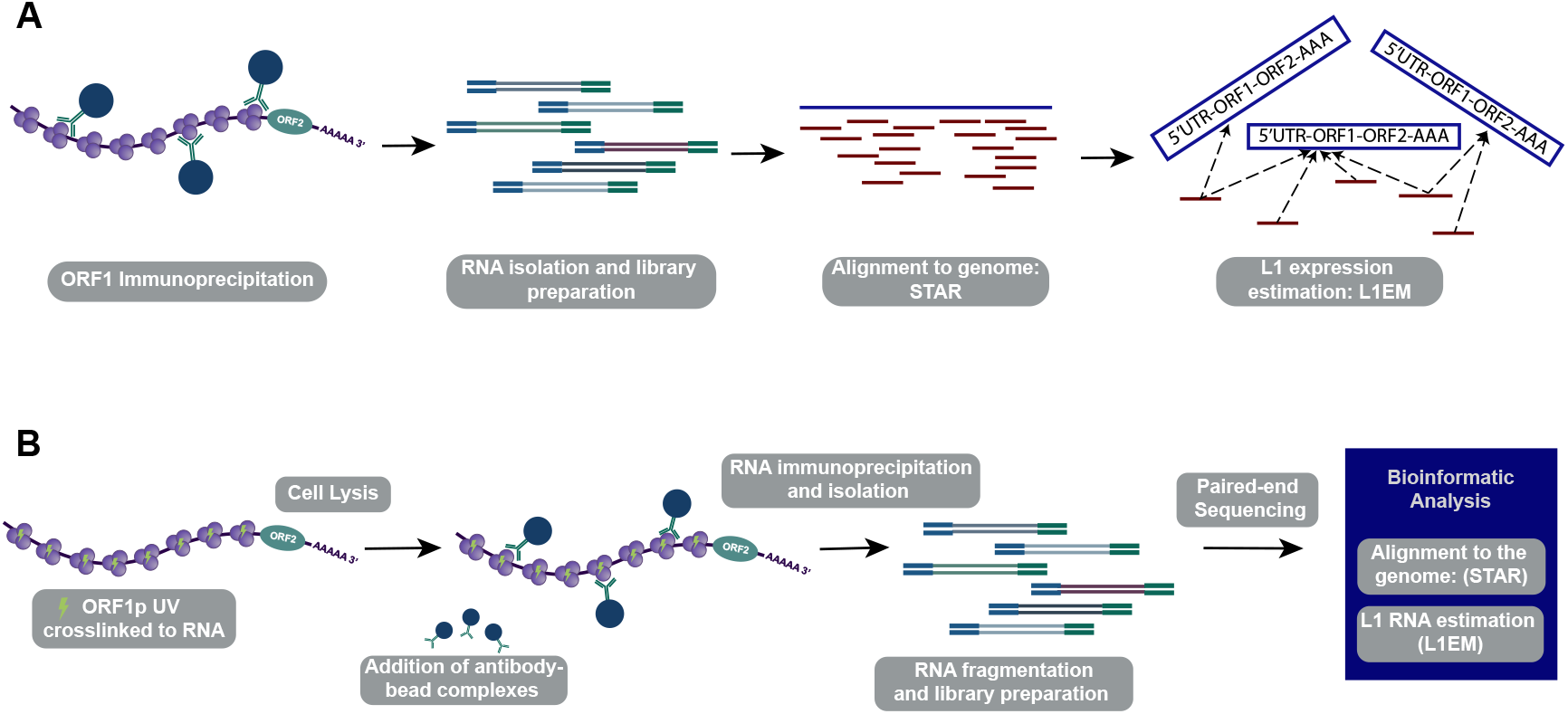
ORF1p RNA Immunoprecipitation. (A) Initial RIP-seq experiments without crosslinking. (B) UV Crosslinked RIP-seq work flow.

**Figure 2.**
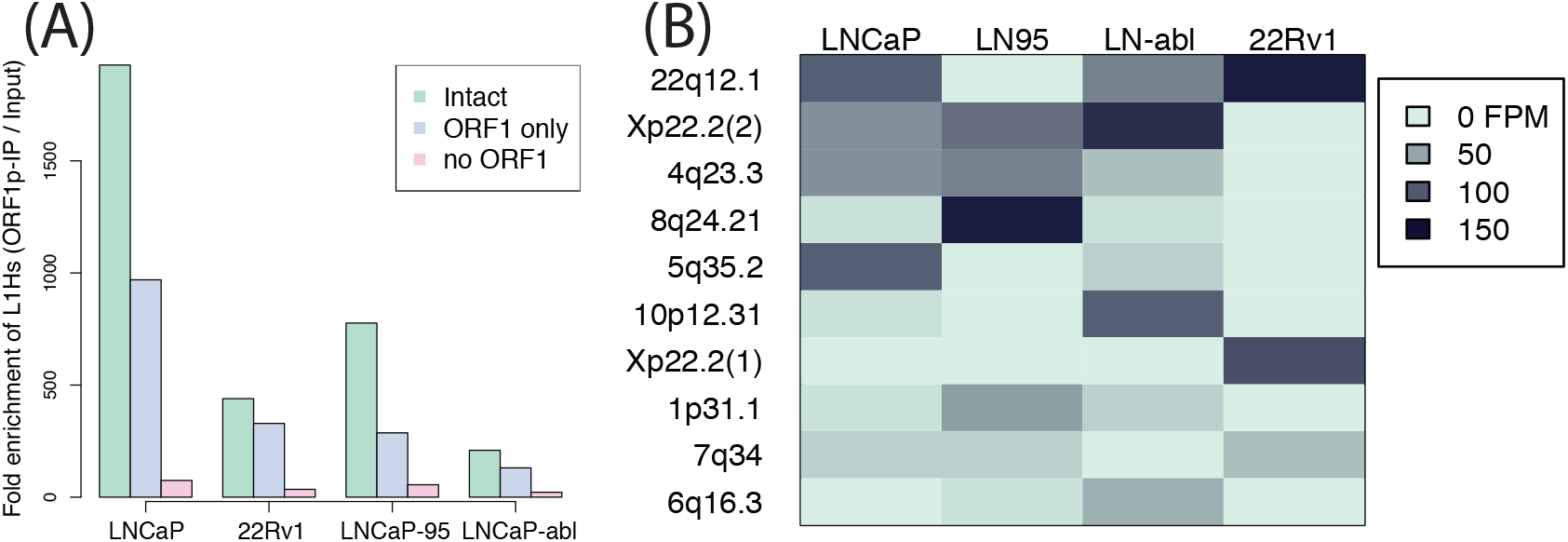
LINE-1 loci present in ORF1p IPs. (A) Total enrichment over input for loci with LINE-1 fully intact (green), with ORF2 truncated, but ORF1 intact (blue) and with ORF1 truncated (pink). Loci with intact ORF1 are more enriched than those with truncated ORF1. (B) Heatmap of the 10 most prevalent intact specific loci IPed in each cell line. 15 additional loci in are shown in S2. 3’ transduction from the highlighted loci have been identified in prostate cancer [28, 65]. It is unclear whether the other expressed loci do not retrotranspose in prostate cancer, have not yet been identified due to limited analyses, or jump without forming 3’ transductions.

### ORF1p IPs are enriched for polyA RNAs

While the enrichment for LINE-1 RNAs with intact ORF1 indicates that we were successful in immunoprecipitating LINE-1 RNPs, we were surprised by the fact that LINE-1 RNA was only a small fraction of the immunoprecipitated transcripts. Previous LINE-1 overexpression studies have shown that ORF1 bound mRNA consisted of 8.3%– 10.3% LINE-1 mRNA, yet our findings examining endogenous ORF1p found even lower levels of ORF1p bound LINE-1 [1]. Reads from younger LINE-1 families (L1Hs, L1PA2, L1PA3, L1PA4) ranged from 0.1% of all reads in LNCaP to 0.15% of all reads in LNCaP-abl. We therefore wondered whether LINE-1 mRNA was dissociating from ORF1p during the IP procedure, freeing ORF1p to interact with other mRNA species present in the cell lysates. To test this possibility, we performed UV crosslinking of LNCaP cells prior to a 1-hour or 3-hour immunoprecipitation. These IPs yielded slightly less LINE-1 RNA: <0.1% of reads in all experiments, indicating that ORF1p was unlikely to disassociate from the LINE-1 mRNA during the IP and that most of the identified interactions of ORF1p and LINE-1 mRNAs formed in the cells, before lysis. Analysis of the non-LINE-1 RNA present in the ORF1p immunoprecipitates showed that, in all experiments, the majority of reads represented ribosomal RNA, which was not depleted from the pool of mRNAs in our experimental design. However, we also found a much greater fraction of reads aligning to exonic sequences in ORF1p IP compared to control (input or IgG IP) (Figure 3A), suggesting an overall enrichment for mature (spliced) gene transcripts in the ORF1p IP. We then calculated the ORF1p co-IP to input RNA ratio for all genes in all four cell lines, finding that most genes are enriched in the ORF1p IP. The median gene enrichment ranged from 5-fold for 22Rv1 to 22-fold for LNCaP (all without crosslinking) (Figure 3B-E). However, in all cases, the enrichment for LINE-1 RNA with intact ORF1 (ranging from 51-fold enrichment over input samples for LNCaP-abl cells to 331 fold enrichment for LNCaP) eclipses the enrichments of all but a few dozen genes, In LNCaP there were 10 genes whose enrichment exceed LINE-1, in LNCaP-95 there were 20, 68 in LNCaP-abl and 24 in 22Rv1. While we did find certain classes of RNA to be enriched by ORF1p IP (see below), we did not identify RNAs that were consistently enriched more strongly than LINE-1. 7 mRNAs did exceed LINE-1 RNA enrichment values in two of the four considered cell lines: *HLTF*, *PCDHGA9*, *PCDHGB5*, *RELN*, *SMC2*, *SI*, and *SP110*. The full list of mRNAs enriched in each experiment can be found in table S2.

**Figure 3.**
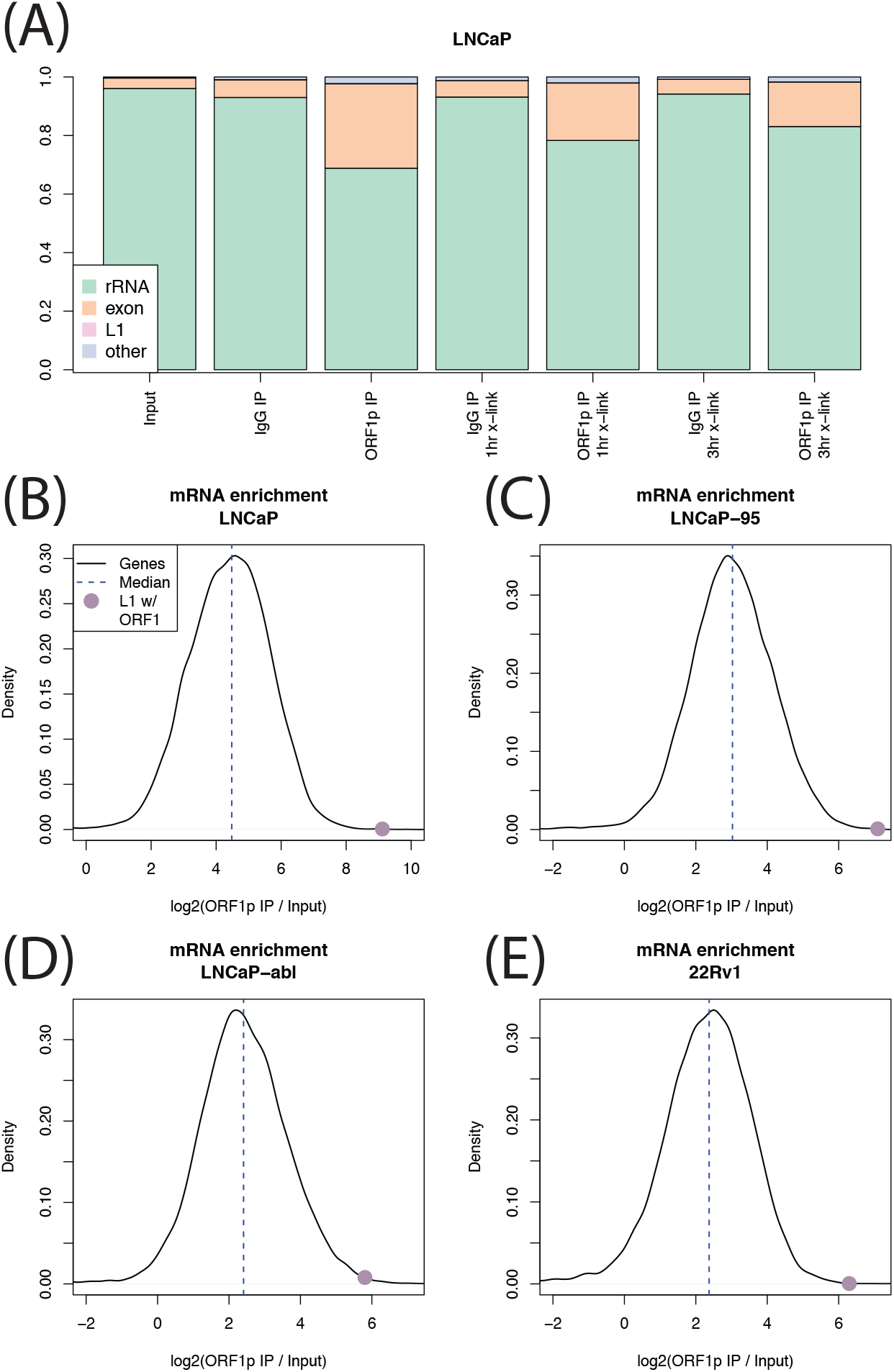
Exonic sequences enriched in IPs. (A) Relative fraction of rRNA, exon aligned RNA, L1 RNA and other RNA in each LNCaP experiment. In all experiments, L1 abundance is too low to be seen. (B-E) Black lines show the distribution of IP enrichment across all genes in each cell line. Purple dot shows LINE-1 mRNA enrichment. In all cell lines, most genes are enriched, but ORF1 is among the strongest enrichments.

The global enrichment for exonic sequences led us to hypothesize that ORF1p is promiscuously binding polyadenylated RNAs. We therefore looked at histone mRNAs, which rely on a unique expression mechanism and are not polyadenylated, expecting a low enrichment for these transcripts in our ORF1p IP samples. We found this to be the case in the LNCaP crosslinking experiments (Figure 4 B-C, mean enrichment 1.1x), but not in the experiments without crosslinking (Figure 4A). This discrepancy indicates that, without crosslinking, there may be some RNAs that do in fact exchange during the IP. Interestingly, as noted above, this does not appear to be the case for LINE-1 RNA itself, enrichment of which was not increased by crosslinking. The consistency of LINE-1 transcripts pulled down in cross-linked and non-cross-linked ORF1 RIPs may be due to ORF1p binding LINE-1 RNA with a higher affinity. However, we cannot exclude the possibility that the high number of ORF1p trimers bound to each LINE-1 mRNA transcript may impact the effect of crosslinking on the bound RNAs.

**Figure 4.**
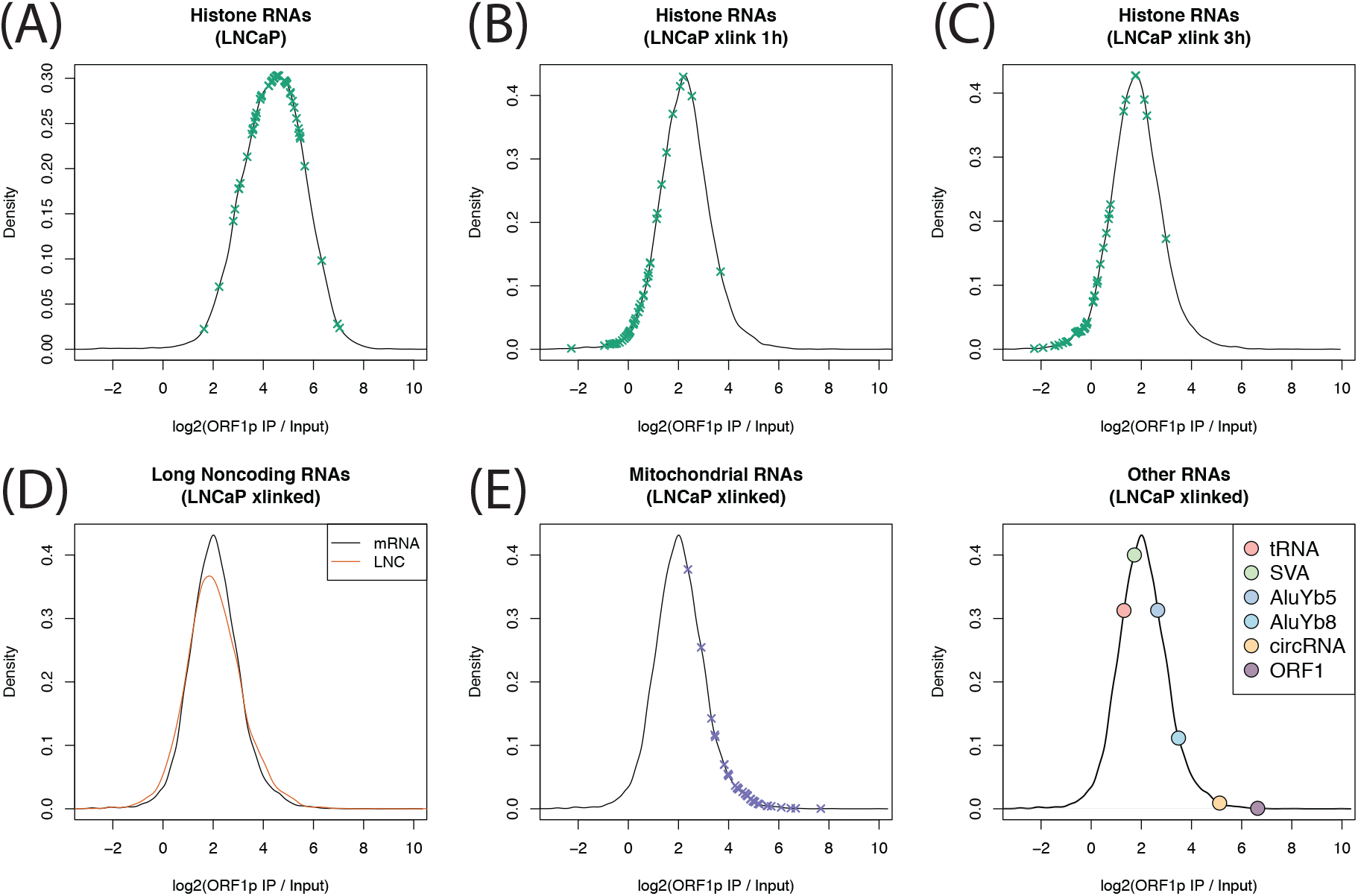
Gene set enrichments. Black lines are the distribution of enrichment across all genes. X’s indicate enrichments in a particular category. Without crosslinking (A), enrichment of histone genes is similar to the enrichment of other genes, but with crosslinking (B, C), enrichment of histone genes is much less. (D) Enrichment is similar for genes that do and do not encode protein. (E) Mitochondrial RNAs are highly enriched in ORF1p IPs. (F) Enrichment of other RNA species. tRNAs are less enriched that most genes, but circRNAs are highly enriched.

We then wanted to know whether the IP enrichment is specific to protein coding mRNAs. lncRNAs, which are polyadenylated, but not translated, showed similar enrichment to mRNA sequences. Median mRNA enrichment was 2.05 fold over input, and median lncRNA enrichment was 2.03x (Wilcox p=0.09, Figure 4D). We also found a strong and surprising enrichment for mitochondrial genome encoded RNA (MT-RNA) in the crosslinking experiments (Figure 4E). This result was unexpected as ORF1p has not been observed to localize inside of mitochondria, but might reflect an association between mitochondria and p-bodies or a colocalization of ORF1p with RNA released from dysfunctional mitochondria presumably into p-bodies (see discussion). Finally, we calculated enrichment for other RNA species, including tRNA, active non-autonomous retrotransposons (SVA, AluYb5, AluYa8) and circular RNAs (circRNA), (Figure 4F). Of these, only circRNA showed strong enrichment (34x enrichment on average). AluYb8 also showed some enrichment (8x on average). No individual circRNA was supported by 10 or more reads, but total circRNA was strongly enriched in the ORF1p IPs. This result likely reflects the known fact the circRNAs locate to p-bodies in the cytoplasm [42] (see next section).

Finally, we asked whether particular classes of mRNA are specifically enriched in ORF1p IPs. We chose to focus on the crosslinking experiments as those appeared more robust (see above). We therefore performed Gene Set Enrichment Analysis (GSEA) [43] using Reactome [44], KEGG [45] and GO [46] gene sets. For all three gene sets, the most highly enriched category contained a large number of ribosomal proteins: ribosome (KEGG), translation initiation (Reactome), and cytosolic ribosome (GO). Ribosomal protein RNAs are some of the most highly expressed mRNAs and are frequently processed into pseudogenes [47], which, given the involvement of ORF1p in retrotransposition of pseudogenes, indicates that these mRNAs can bind directly to ORF1p. Overall, there is a weak correlation between the number of times a gene has been processed into a pseudogene [48] and enrichment in the ORF1p IP (Spearman *ρ* = 0.11, *p* = 1.1 × 10^−6^). Only a few of the gene sets enriched at FDR<5% are not dominated by ribosomal proteins. These are: propanoate metabolism (KEGG), apoptotic cleavage of cellular proteins (Reactome), apoptotic activation phase (Reactome), cilium or flagellum dependent cell motility (GO), and homophilic cell adhesion via plasma membrane adhesion molecules (GO). Enrichment in this last category is driven by protocadherins.

### ORF1p associated mRNA is enriched in p-body RNA

We reasoned that ORF1p localization to some phase separated RNP granules may explain why the RNAs that IPed with ORF1p were particularly enriched for polyA RNAs. Because ORF1p is primarily cytoplasmic [7], two types of granules are natural candidates: stress granules (SG) and processing bodies (p-bodies). We therefore compared our ORF1p enrichments from the LNCaP crosslinking experiments to published SG [49] and p-body RNA enrichments [50]. These comparisons are not perfect as the experiments were not done in LNCaP cells, but they can still provide a sense of whether the IPed mRNAs are SG- or p-body-like. We found a strong correlation between the genes enriched by ORF1p IP and those enriched by p-body purification (Spearman *ρ* = 0.4, *p* ≈ 0, Figure 5A), with more than half (63%) of the genes that are significantly (BH FDR<5%) enriched in the ORF1p IP also enriched in the p-bodies (odds ratio = 2.5, *p* = 2.6 × 10^−77^, Figure S3A). There was also a correlation with SG enrichment (Spearman *ρ* = 0.15, *p* = 7.1 × 10^−43^, Figure 5B) and a significant overlap between genes enriched in ORF1p IP and those enriched in SG (odds ratio =1.3, *p* = 1.1 × 10^−5^, Figure S3B), but the correlation becomes negative when using partial correlation to account for the similarity between p-bodies and SGs (*ρ* = −0.08, *p* = 1.4 × 10^−12^), indicating that the positive relationship between ORF1p bound and SG localized RNA can be explained by the fact that many RNAs are enriched in both p-bodies and SG. This result indicates that ORF1p interacts with mRNAs that locate to p-bodies.

**Figure 5.**
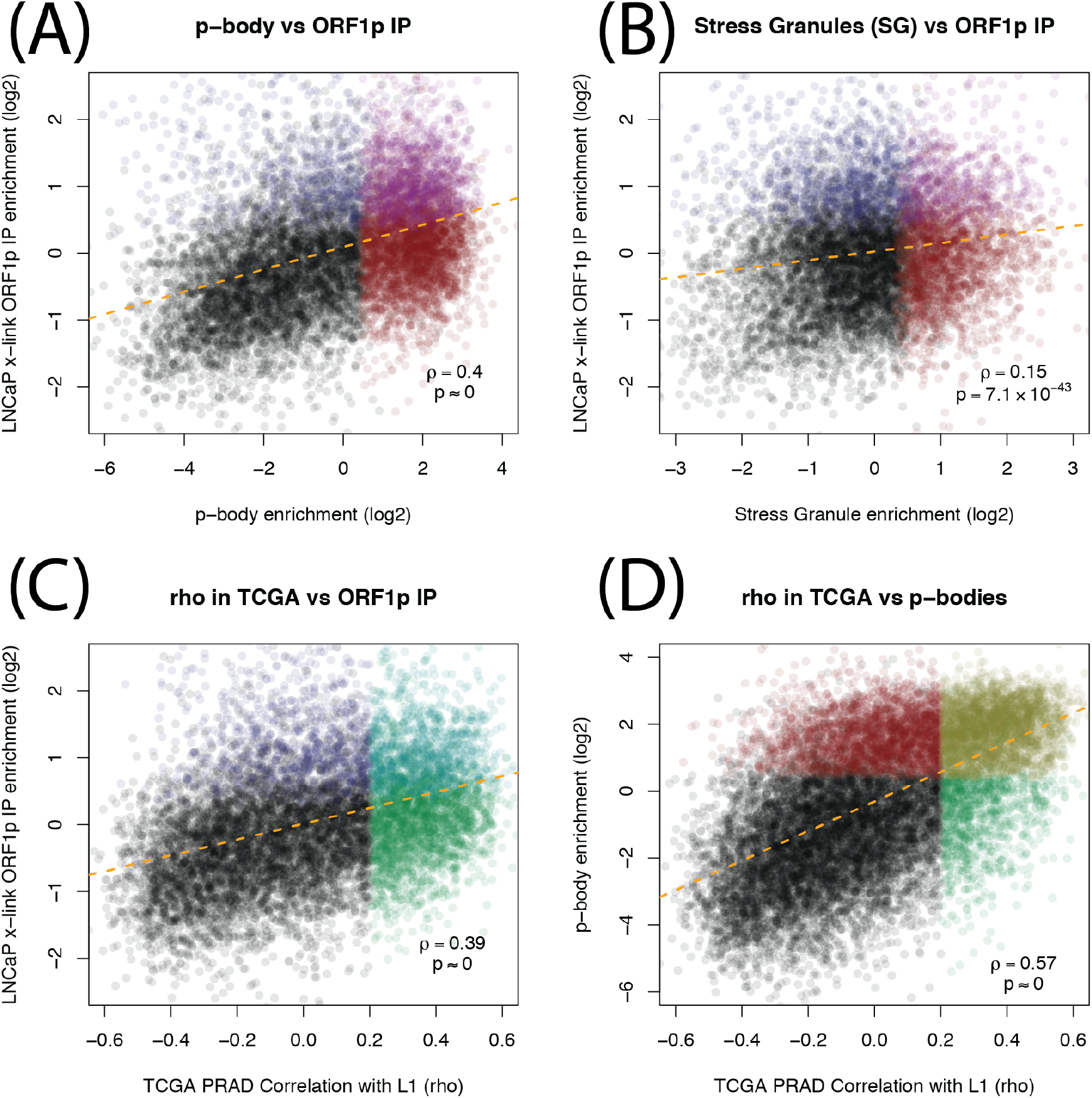
Scatter plots comparing ORF1p IP RNAs, granules RNAs and RNAs correlated with LINE-1 in TCGA prostate cancer. (A) For each gene, enrichment in p-body purification study [50] (x-axis) vs enrichment in our ORF1p IPs, calculated by DESeq2 (y-axis). (B) Same y-axis as in A, but using SG enrichments [49] on the x-axis. (C) RNA expression correlation between host genes and intact LINE-1 in TCGA prostate cancer samples (x-axis) vs enrichment in our ORF1p IPs (y-axis). (D) Same x-axis as in C, but y-axis is p-body enrichment (same as x-axis in A).

Because p-bodies are important regulators of mRNA processing, we hypothesized that LINE-1, or at least ORF1p expression, could have a wide impact on the abundance of certain mRNAs. To investigate this, we used L1EM to quantify LINE-1 RNA in prostate tumor RNA-seq available through TCGA. We found that genes significantly correlated with LINE-1 expression were more likely to be enriched in the ORF1p IP (odds ratio = 2.5, *p* = 1.2 × 10^−73^, Figure S3C), and that there is a positive relationship between correlation with LINE-1 RNA in TCGA and enrichment in our ORF1p IP experiments (*ρ* = 0.39, *p* ≈ 0, Figure 5C). More strikingly, most (72%) of the genes that were significantly correlated with LINE-1 expression were also enriched in p-bodies (odds ratio = 7.5, *p* ≈ 0, Figure S3D), and there is an even stronger positive relationship between LINE-1 RNA expression in TCGA and enrichment in p-bodies.

## Discussion

The reactivation of LINE-1 expression in prostate cancer has raised new questions regarding its role in cancer initiation and progression. Sequencing studies have shown active retrotransposition in prostate tumors, and an increased rate of LINE-1 retrotransposition in metastatic prostate cancer compared to primary tumors. However, the expression profiles of LINE-1 loci (mRNA) during prostate cancer progression has not been widely examined. We have previously shown an increase in ORF1p expression in androgen independent LNCaP cell lines, LNCaP-95 and LNCaP-abl, compared to parental LNCaP cells [21]. Here we show an overall increase in LINE-1 mRNA expression in both LNCaP-abl and LNCaP-95 cells compared to the parental LNCaP line. While LNCaP related cell lines (LNCaP, LNCaP-abl, LNCaP-95) show stronger similarities in LINE-1 loci expression when compared to non-related cell lines (22Rv1), the profile of LINE-1 loci changes significantly in the androgen independent clones compared to androgen sensitive cells. Together, these findings may offer insight into the increased rate of retrotransposition in metastatic prostate cancer. Additionally, several of the loci expressed in these cell lines have generated new somatic insertions previously identified in actual human prostate tumors, indicating that at least some of the expressed loci are actually capable of retrotransposition.

Unexpectedly, we found that among all the RNA bound by ORF1p only a small percentage is represented by LINE-1 mRNA. While our analysis showed that our ORF1p IP was enriched for full length and complete ORF1p LINE-1 mRNA, indicating successful LINE-1 RNP immunoprecipitation, the majority of RNA enriched in our ORF1 IP consisted of non-LINE-1 transcripts (figure 6). Interestingly, mature exonic mRNAs were highly enriched in the ORF1-IP, indicating an enrichment of polyadenylated RNAs and generally speaking cytoplasmic as opposed to nuclear location. The lower levels of the non-polyadenylated histone mRNA in the ORF1 IP confirmed ORF1p’s propensity to bind polyA mRNA. ORF1p has been shown to bind nucleic acids in a sequence independent manner [16]. These findings make it unlikely that ORF1p is binding polyA sequences due to a sequence preference. ORF1p may also associate with polyA mRNA through its interaction with other proteins. For example, polyadenylate-binding protein 1 (PABPC1), a polyA binding protein necessary for efficient retrotransposition, has also been shown to interact with ORF1p [51]. This interaction, as well as ORF1p’s localization in the cytoplasm, may all increase its tendency to bind polyA mRNAs.

**Figure 6.**
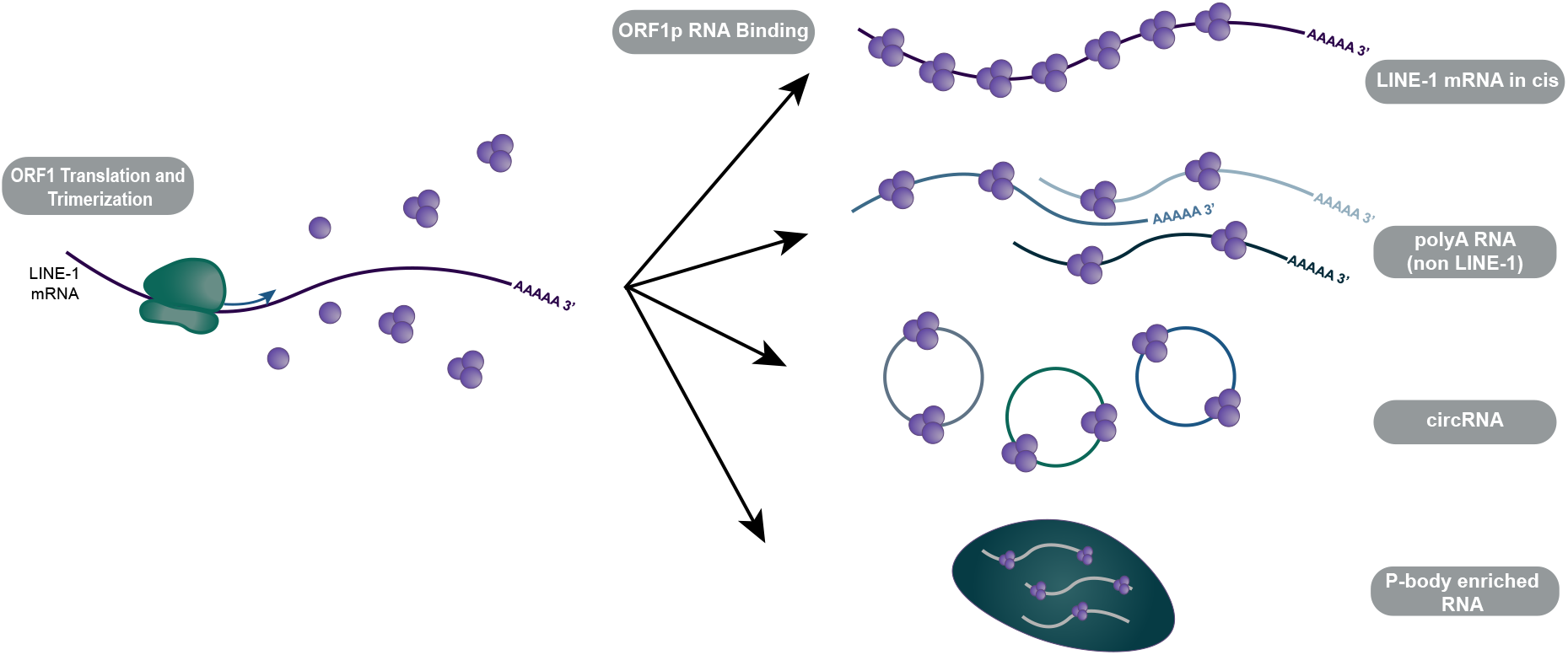
RNAs enriched in ORF1p IPs. LINE-1 mRNA with full length ORF1 are most strongly enriched. A wide range of mRNAs and circRNAs are also enriched in ORF1p IP. Most of the significantly enriched mRNAs were also enriched in a previous study that isolated RNA from p-bodies. Consistent with this, circRNA are also believed to located to p-bodies.

We were particularly surprised to find MT(mitochondrial DNA encoded)-RNA enrichment in the crosslinked ORF1p IP experiments. MT-mRNAs are polyadenylated, and it is possible that ORF1p could bind them directly. While no study has shown ORF1p localizing in the mitochondria, ORF1p may bind MT-RNA that has been release from dysfunctional mitochondria. However, we did also find that MT-rRNAs are also enriched (although to a lesser degree). We considered whether this enrichment might be the result of MT-DNA contamination as mtDNA contamination can be amplified by the many MT genome copies that exist in each cell. Globally, we find that the vast majority of reads align to rRNA or exons, indicating that DNA contamination is not likely to be a significant factor in our analysis. However, MT specific DNA contamination is difficult to rule as most of MT genome is transcribed as a single transcript. A third possibility is that crosslinking connects mitochondria to nearby ORF1p containing RNPs. Mitochondria are involved in interferon signaling [52] and RNA interference [53], both of which are involved in LINE-1 repression [54], and p-bodies have been shown to locate near mitochondria [53]. It is feasible that cytoplasmic ORF1p and LINE-1 RNPs may localize in close proximity to mitochondria and can be connected by crosslinking, leading to the extraction of entire mitochondria during the IP step.

Given strong evidence from previous studies indicating that ORF1p locates to stress granules (SG), we expected to find a large overlap between ORF1p bound mRNAs and SG enriched mRNAs. Instead, we found only a small overlap that can be explained by the overlap between p-body and SG RNA. In contrast, we found a much larger overlap between ORF1p bound mRNAs and p-body enriched mRNAs compared to SG RNAs. This may indicate that endogenously expressed ORF1p localizes to p-bodies in LNCaP cells, or it may indicate that ORF1p promotes the mislocalization of p-body RNAs to sites of ORF1p accumulation. Alternatively, the high levels of L1 sequences may lead to the formation of a third type of body/granule with intermediate properties.

The discrepancy between our results and the previous studies pointing to SG as sites of LINE-1 proteins/RNA localization, may be explained by the fact that LNCaP cells were “unstressed” in our study. Goodier et al. (2007) did find endogenous ORF1p localizing with SG markers in embryonal carcinoma cell lines [20]. However, this localization was most pronounced after exogenous stress. It may be that LNCaP cells are able to tolerate endogenous LINE-1 expression with minimal stress, leading to a lack of SG for ORF1p to localize to. Another major difference between our study and this previous study is that our study used RNA sequencing – a much more sensitive assay compared to the immunofluorescence and microscopy approaches used by Goodier et al. Overall, LINE-1 may localize to SG or to SG-like granules, but, at least in LNCaP cells, the mRNAs in those granules may be more similar to p-body mRNAs than to the mRNAs present in canonical arsenite-induced SGs.

Whether ORF1p localizes to p-bodies or causes p-body RNA mislocalization, it may be interfering with the normal processing of p-body RNAs. Our results show that ORF1p-bound mRNAs expression correlates with LINE-1 expression in prostate cancer. Thus, LINE-1 RNPs (at least the ones containing ORF1p) may be interfering with the processing/degradation of certain mRNAs. In particular, almost all of the genes that are significantly correlated with LINE-1 RNA in prostate cancer were also enriched in p-bodies, indicating that LINE-1 may interfere with the degradation of p-body associated RNAs. In yeast, p-body proteins are involved in the regulation of gene expression related to DNA replication stress resistance [55], possibly indicating a link between our findings and the previously documented relationship between LINE-1 and replication stress [36, 56].

## Conclusions

Our study finds that the increased LINE-1 expression in the androgen independent LNCaP-95 and LNCaP-abl cells over the parental LNCaP is accompanied by a large-scale remodeling of the expressed LINE-1 loci. We also find that ORF1p associates not only with LINE-1 mRNA but also with a wide range of non-LINE-1 transcripts, particularly polyA RNAs. ORF1p bound RNA transcripts are enriched for p-bodies localized RNAs. Notably, the ORF1p-bound and p-bodies localized RNA species also correlate with LINE-1 expression in prostate cancer raising the intriguing possibility that cytoplasmic ORF1p may affect RNA processing in prostate cancer cells.

## Methods

### Cell Culture

LNCaP (CRL-1740) and 22Rv1 (CRL-2505) cells were purchased from ATCC and maintained in RPMI 1640 with 10% FBS. LNCaP-abl, and LNCaP-95 cell lines were generous gifts from Z. Culig, and J. Isaacs, respectively and maintained in RPMI 1640, phenol red free, and 10% charcoal dextran stripped FBS. Cells were regularly screened for mycoplasma.

### RNA Immunoprecipitation

For crosslinked samples, cells were kept on ice, washed twice with cold PBS, and UV-crosslinked using a Stratalinker 2400 with 150 mJ/cm^2 at 254nm. Four 15cm plates of each cell type, ~70% confluency, were used for each immunoprecipitation. RNA immunoprecipitation was conducted using the Magna RIP RNA-Binding Protein Immunoprecipitation Kit (Millipore Sigma 17-700) according to manufacturer’s protocol. Magnetic beads were incubated with 10μg ORF1p (Millipore Sigma MABC1152) or mouse IgG (Santa Cruz sc-2025) antibody per 50μl of magnetic beads. Bead/antibody complexes and cell lysates were incubated for either 1hr or 3hrs at 4°C. RNA was isolated using a phenol chloroform extraction, following the suggested protocol from the Magna RIP kit, and an on-column DNase digestion and RNA cleanup was performed using a RNeasy MinElute Cleanup Kit (Qiagen 74204, 79254).

### RNA Library Preparation and Sequencing

Sequencing library was prepared using the NEBNext Ultra II RNA Library Prep Kit for Illumina (NEB E7770S) according to manufacturer’s protocol, each sample prepared with its own unique barcode (NEB E7600S). Prepared libraries were sequenced as paired end 36-cycle reads (20M) on the NextSeq 500. Reads were demultiplexed with Illumina bcl2fastq v2.20 requiring a perfect match to indexing BC sequences.

### Alignment and LINE-1 RNA identification

Reads were aligned to the hg38 human reference genome using the STAR aligner and TCGA mRNA analysis pipeline options [57]. Locus specific LINE-1 RNA expression was estimated using L1EM [41]. ORFs >300aa were translated from Repeatmasker annotated LINE-1 (L1Hs, L1PA2, L1PA3, L1PA4) sequences in hg38 using ORFfinder (NCBI) and then aligned to ORF1p and ORF2p consensus sequences from Dfam using BLAST [58]. ORFs were considered intact if the alignment covered at least 95% of the consensus sequence. Intact LINE-1 and ORF1 expression were calculated by adding the estimated expression for all loci with intact ORFs.

### Quantification of non-LINE-1 RNA

rRNA was quantified using samtools [59] to count reads overlapping rRNA genes annotated in the UCSC genome browser repeat track [60]. These reads were then filtered out and reads overlapping UCSC genome browser annotated exons were counted by samtools. Other repetitive RNAs were quantified using bedtools and the UCSC genome browser repeat track [60]. Circular RNAs were identified and quantified using CIRIquant [61]. Because no circRNA was supported by a large number of reads, all circRNA reads were pooled into a single quantification. Reads aligning to each gene in GRCh38.96 were counted using featureCounts [62]. The LNCipedia database was used with featureCounts to quantify lncRNA reads [62].

### ORF1p IP, p-body, SG and TCGA comparisons

ORF1p IP enrichment for each gene was estimated using DEseq2 [63]. Note that DESeq2 normalizes expression to exon aligned reads, rather than all aligned reads leading to the difference in scale for figure 5 compared to figures 3 and 4. P-body enrichments were obtained from Hubstenberger et al. [50] and SG enrichments were obtained from Khong et al. [49] LINE-1 expression was estimated in TCGA prostate cancers using L1EM as implemented on the Cancer Genome Cloud (CGC) [64]. Spearman correlation and significance between LINE-1 RNA and upper quantile normalized expression for each gene (available from the Genomic Data Commons (GDC)) was calculated using in R.

## ABREVIATIONS

LINE-1: Long Interspersed Element-1
L1EM: LINE-1 Expectation Maximization
SG: Stress granule
P-body: Processing body
IP: Immunoprecipitation
RNP: ribonucleoprotein

## DECLARATIONS

## Acknowledgements

Sequencing was performed by the NYU Langone Institute for Systems Genetics. We would like to thank Megan Hogan and Raven Luther for their technical assistance and expertise.

## Funding

This work was supported by the National Institutes of Health Grants R01CA112226 (to S. K. L.), 1F31CA225053-01A1 (to E.M.B.) and P01AG051449 (subcontract to J.D.B. and D.F.). This project has been funded in part with Federal funds from the National Cancer Institute, National Institutes of Health, under Contract No. HHSN261200800001E (subcontract to W.M.).

## Conflict of Interest

The authors declare that they have no conflicts of interest with the contents of this article. The content is solely the responsibility of the authors and does not necessarily represent the official views of the National Institutes of Health.

## Author Contributions

Experiments were performed by EMB with help from PM. Data and statistical analyses were done by WM and DF. Experiments were planned and analyzed by EMB, WM, PM, SKL, and DF. Experiments were supervised by SL, JDB and DF. WM, EMW, SKL and DF wrote the manuscript.

## Competing Interests

The authors declare that they have no competing interests

## Availability of Data and Material

Raw Illumina sequencing reads are available from the SRA database under bioproject XXXXXXXX.

## Consent for Publication

Not applicable

## Ethics Approval and Consent to Participate

Not applicable

---

**Figure S1.**
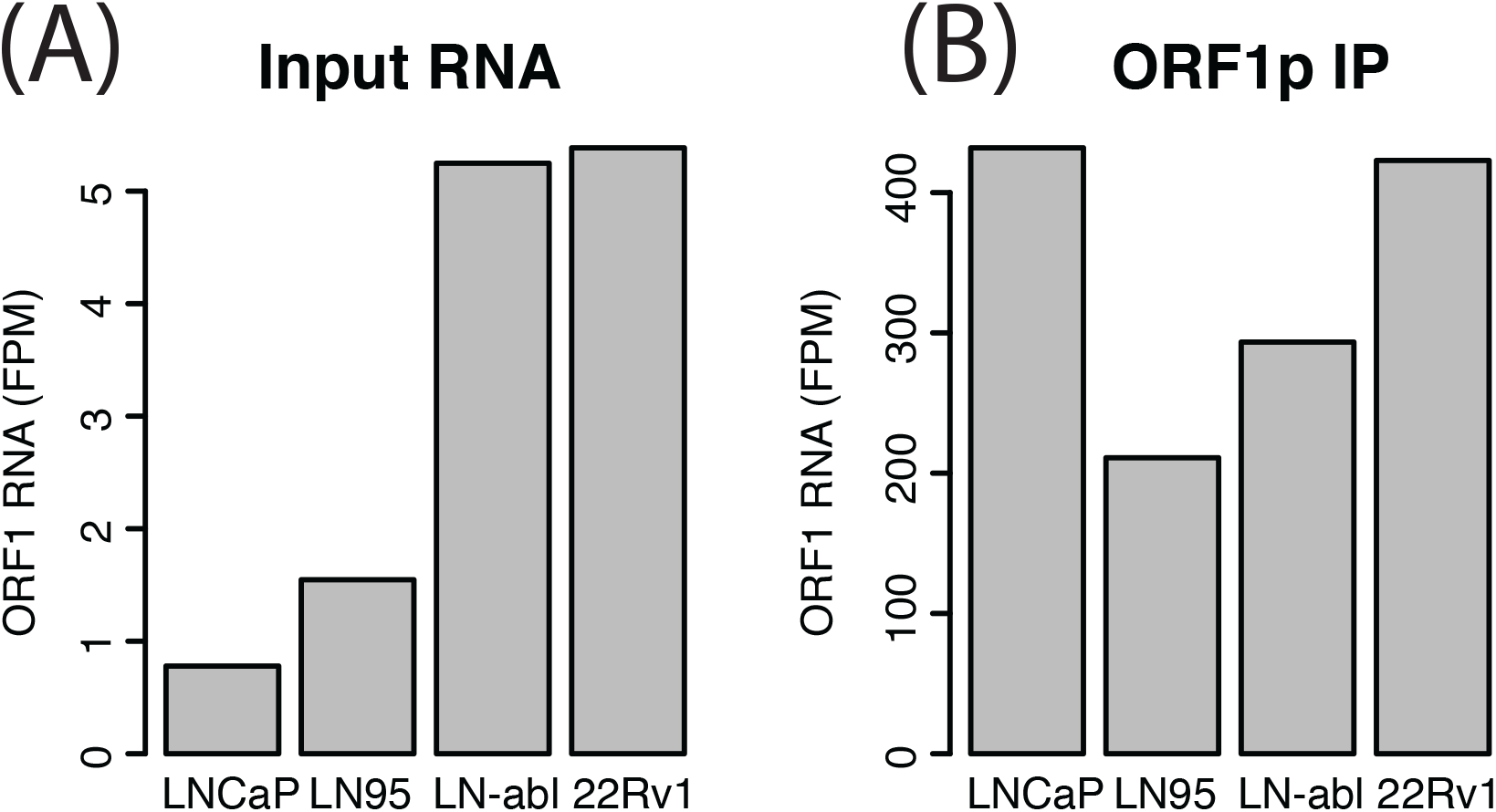
(A) LINE-1 RNA in the input for each cell LINE-1. (B) LINE-1 RNA after ORF1p-IP enrichment.

**Figure S2.**
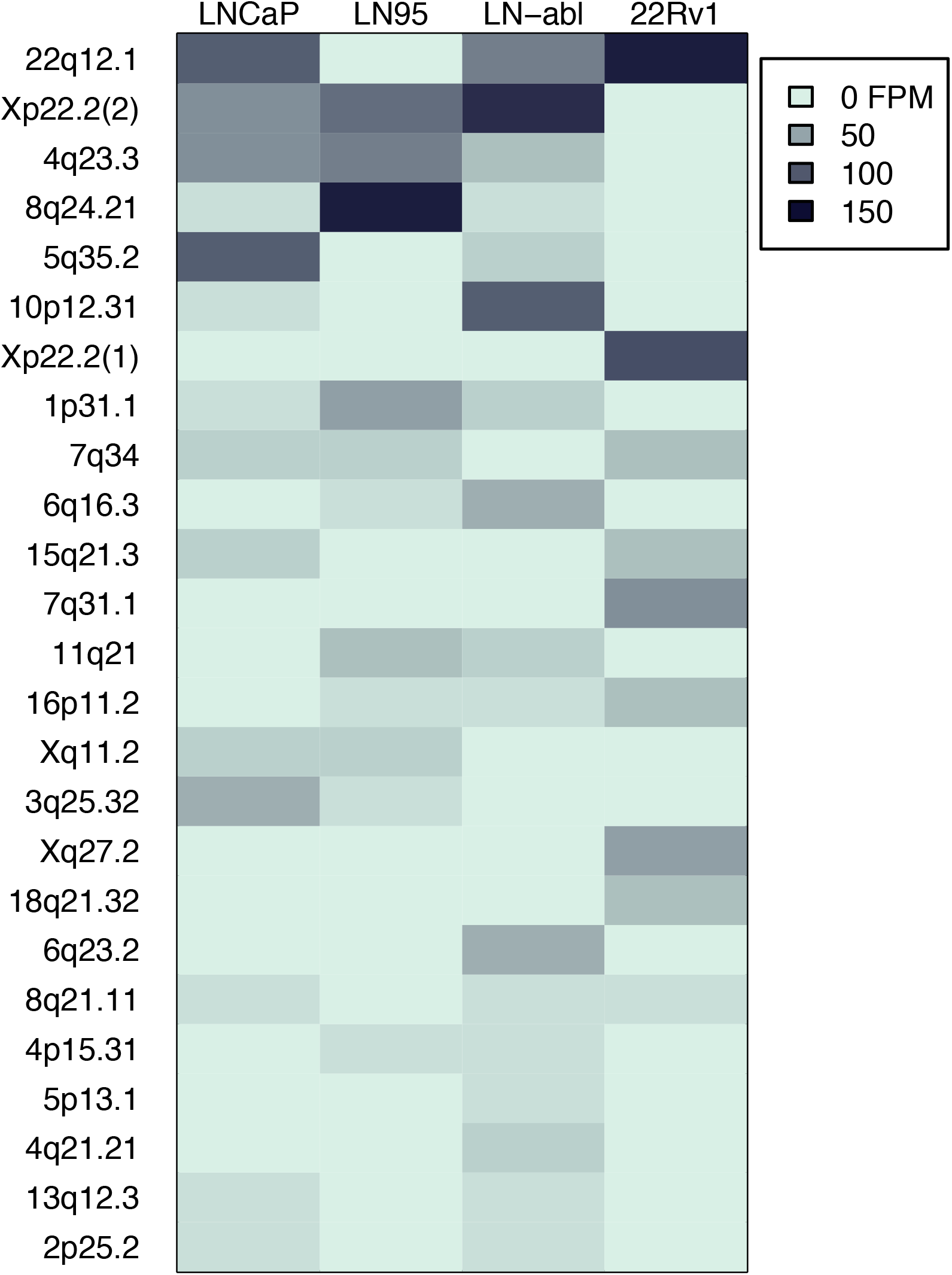
Heatmap of the specific intact loci IPed in each cell line. Expansion of figure 1B to include the top 25 loci.

**Figure S3.**
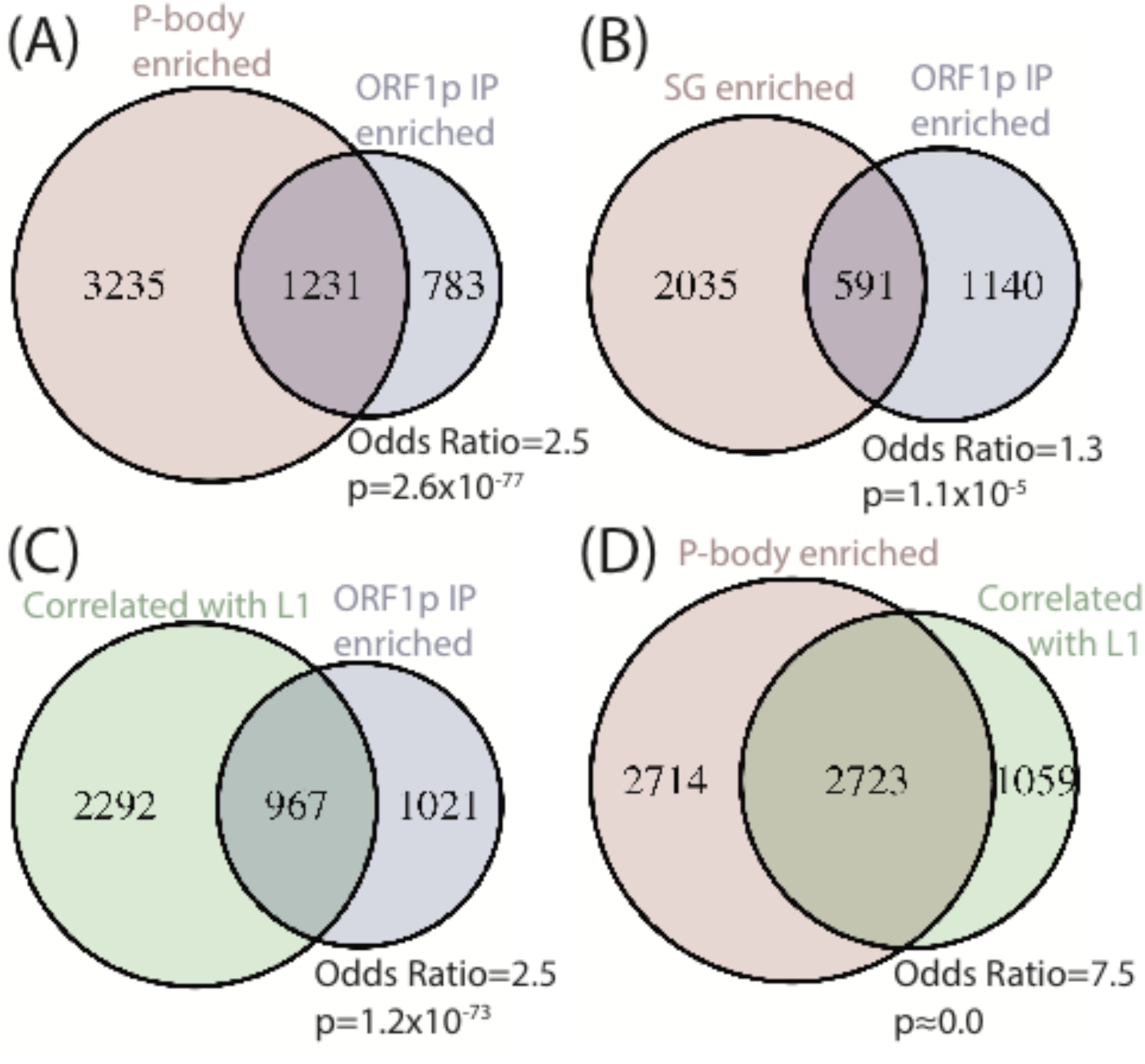
Overlap between transcripts enriched in ORF1p IP and in cytoplasmic granules. (A) Overlap between ORF1p IP enrichment and p-body enrichment [50]. A 5% FDR cutoff was used. (B) As in A, but comparing ORF1p IP to SG enrichment [49]. (C) Overlap between transcripts enriched in ORF1p IP and those whose expression is positively correlated with LINE-1 RNA in TCGA prostate cancer samples (i.e. these genes are more highly expressed in tumors that express more LINE-1). (D) Overlap between transcripts enriched in p-bodies and those correlated with LINE-1 RNA in TCGA prostate cancer.

## Notes

### Competing Interest Statement

The authors have declared no competing interest.

